# Mixed Multi-Level Visual, Reward, and Motor Signals in Dorsomedial Frontal Cortex Area F7 during Active Naturalistic Video Exploration

**DOI:** 10.1101/2023.09.25.559420

**Authors:** Farid Aboharb, Stephen Serene, Julia Sliwa, Winrich Freiwald

## Abstract

In the primate brain the frontal lobes support complex functions, including social cognition. Understanding the functional organization of these regions requires an approach for rich functional characterization. Here we used a novel paradigm, the visual exploration of dynamic social and non-social video scenes, to characterize diverse functions in area F7 of dorsomedial premotor cortex in the macaque monkey (*Macaca mulatta*) previously suggested to be involved in the representation of social interactions. We found that neural populations within this area carry information about both visual events in the videos, like head turning, and higher-level social categories, like grooming. In addition to signaling visual events, the population also encoded the delivery of juice reward. Our novel free viewing paradigm and naturalistic stimuli elicited active visual exploration, and we found that a large fraction of F7 neurons responded to the subject’s own saccadic eye movements. Information from these three different domains were not separated across distinct neural sub-populations, but distributed, such that many neurons carried sensory, reward, and motor information in a mixed format. Thus we uncover a hitherto unappreciated diversity of functions in region F7 within dorsomedial frontal cortex.

## Introduction

Understanding the neural mechanisms of complex functions requires identifying the relevant brain areas and dissecting the specific sub-functions each area supports. A case in point are recent successes in dissecting the neural mechanisms of face recognition. The localization of face-selective regions in the macaque monkey brain with functional magnetic resonance imaging (fMRI)(Tsao et al., 2003); the demonstration, with single-cell electrophysiology, that these areas are highly specialized for face-processing (Tsao et al., 2006); and demonstrations of their causal roles in face-processing behaviors (A. Afraz et al., 2015; S.-R. Afraz et al., 2006; Moeller et al., 2017; Sadagopan et al., 2017) identified the neural substrates of face-processing and thus made it possible to determine the neural mechanisms of a complex function. Towards this goal, systematic analyses of neural activity during the presentation of large image sets, showed that each area generates a unique code for faces (Freiwald et al., 2009), and how these codes could explain major properties of face recognition (Tanaka & Farah, 1993; Taubert et al., 2015; Yin, 1969). Location, connectivity, and functional properties of face areas like response latencies and invariance properties further suggested a specific hierarchical transformation of the codes from one area into that of the next (Farzmahdi et al., 2016). These investigations combined allowed for the generation of computational models replicating the key properties of the system. These successes will likely translate into neighboring domains within the temporal lobe (Bao et al., 2020; Hesse & Tsao, 2020) and thus uncover general mechanisms of object processing.

The core of this successful research program – the characterization of a large brain region with fMRI followed by detailed electrophysiological investigations of single cells within fMRI-identified brain areas – can also be used in other domains. The current paper is the first component of a research program aiming to take this approach from the domain of vision into the domain of cognition, focusing on the domain of social cognition. While face processing is largely supported by areas in the temporal lobe (Moeller et al., 2008), major areas of social cognition reside in the frontal lobe (Báez-Mendoza et al., 2021; Shepherd & Freiwald, 2018; Sliwa & Freiwald, 2017). Frontal lobe regions have been differentiated based on cytoarchitecture (Morecraft et al., 2012), connectivity (Barbas & Pandya, 1989), function in imaging (Sallet et al., 2013) and lesion studies (Gregoriou et al., 2014; Rossi et al., 2007), as well as single unit recordings (Romanski, 2004). There remains a need to integrate across these different levels of description. Critical to the success of such a program is the use of a common paradigms for functional comparisons across areas. This is particularly important for the frontal lobes whose functions are likely more complex and thus more difficult to characterize than those of the temporal lobes.

Such a paradigm should invoke a wide range of cognitive capabilities, yet not require extensive training; be widely usable across species; and ideally utilize some of the principles that revealed the organization of temporal lobe face processing. Videos of real-world social interaction offer a basis for such a paradigm when paired with suitable control stimuli. Real-world social interactions contain rich visual information embedded in shape, color, and motion. These visual scenes include component stimuli like faces and bodies, for which specialized process areas have already been identified (Premereur et al., 2016), including in the frontal lobe (Tsao et al., 2008). Social interactions unfold on multiple timescales: heads may turn towards each other abruptly, or bodies change their motion direction, yet despite these fast changes, the behavioral state of the interaction can remain constant across seconds. And social interactions are not just complex visual stimuli, they automatically invoke social cognition. Observers are compelled to interpret these spatio-temporal patterns of pixels at a high level (Isik et al., 2017) and make inferences about the underlying and unobserved internal states of participants. This paradigm thus automatically invokes cognition, and no training is required. For this reason, these stimuli can also be used across species. Social interaction videos not only invoke cognitive and emotional responses in an observer (Dziura et al., 2021): they also lead to motor responses. Among these motor responses are eye movements scanning the major points of the social scene. These motor responses are automatic and need not be consequential – the social video will unfold regardless.

A recent fMRI study (Sliwa & Freiwald, 2017) developed and used such a 3^rd^-person social interaction video paradigm and successfully localized areas in the frontal lobes specifically involved in their processing. To do so, control stimuli including non-social goal-directed behavior or idling social agents as well as non-social physical interactions were included. Social interaction stimuli engaged multiple regions within medial, dorso-medial, and ventrolateral frontal cortex (Sliwa & Freiwald, 2017). The broad engagement of prefrontal cortex suggests that these stimuli indeed invoked cognition, and the fact that many areas in different parts of the frontal lobe were engaged suggests that the paradigm activated different functions. Some of the areas overlapped with areas involved in high-level social cognition like mental state attribution (Báez-Mendoza et al., 2021), yet this study could not identify specific functional contributions of the individual areas. Unraveling the mechanisms of these capabilities will require temporally resolved electrophysiological measurements targeted to specific brain areas.

Here we adapted this paradigm for electrophysiological recordings in order to establish its usefulness in characterizing a broad range of functions including sensory, cognitive, and motor functions. In selecting a brain region, we took advantage of a classical observation in the organization of the frontal lobes (Schall, 2015): in dorsomedial frontal cortex, there is a hierarchical arrangement of areas along the caudo-rostral axis with stronger motor functions and shorter response latencies at the caudal end and increasingly cognitive functions toward the frontal end. Dorsomedial frontal cortex has frequently been found to be activated in social cognitive tasks (Fletcher et al., 1995; Goel et al., 1995; Ozonoff et al., 1991), and specifically in the fMRI study using the social interaction video paradigm (Iacoboni et al., 2004). We thus selected the most posterior area, area F7, in which such activation was found, reasoning that it would allow us to determine to what extent specific events or general states in the video would be encoded and whether specific functions (like sensory, cognitive, or motor) could be isolated similar to how specific representations of faces are separated from each other in the temporal lobe (Freiwald et al., 2009).

## Results

We recorded during the presentation of sixteen naturalistic videos (Sliwa & Freiwald, 2017) including videos containing conspecifics engaging in social interactions (two videos containing each of: chasing, fighting, grooming, and mounting behaviors, for a total of 8, Fig 1B), two videos containing non-social goal-directed behavior and two videos containing idling conspecifics, as well as two containing object motion, and two landscape videos. Videos were presented for 2,800 ms after an initial 800-950 ms preparatory period during which the subject fixated a centrally located dot (Fig. 1A). Videos could be freely explored (see Methods), and when gaze was maintained inside the area of the stimulus plus a margin, a juice reward was delivered after 180-220ms, followed by a 500ms long inter-trial interval. To quantify gaze behavior, objects in videos were segmented and gaze patterns were analyzed (Fig 1C). Despite no extrinsic incentive, subjects consistently attended to conspecifics in videos, spending most of the time attending to bodies, faces, and hands (Fig 1D).

**Figure 1 -.**
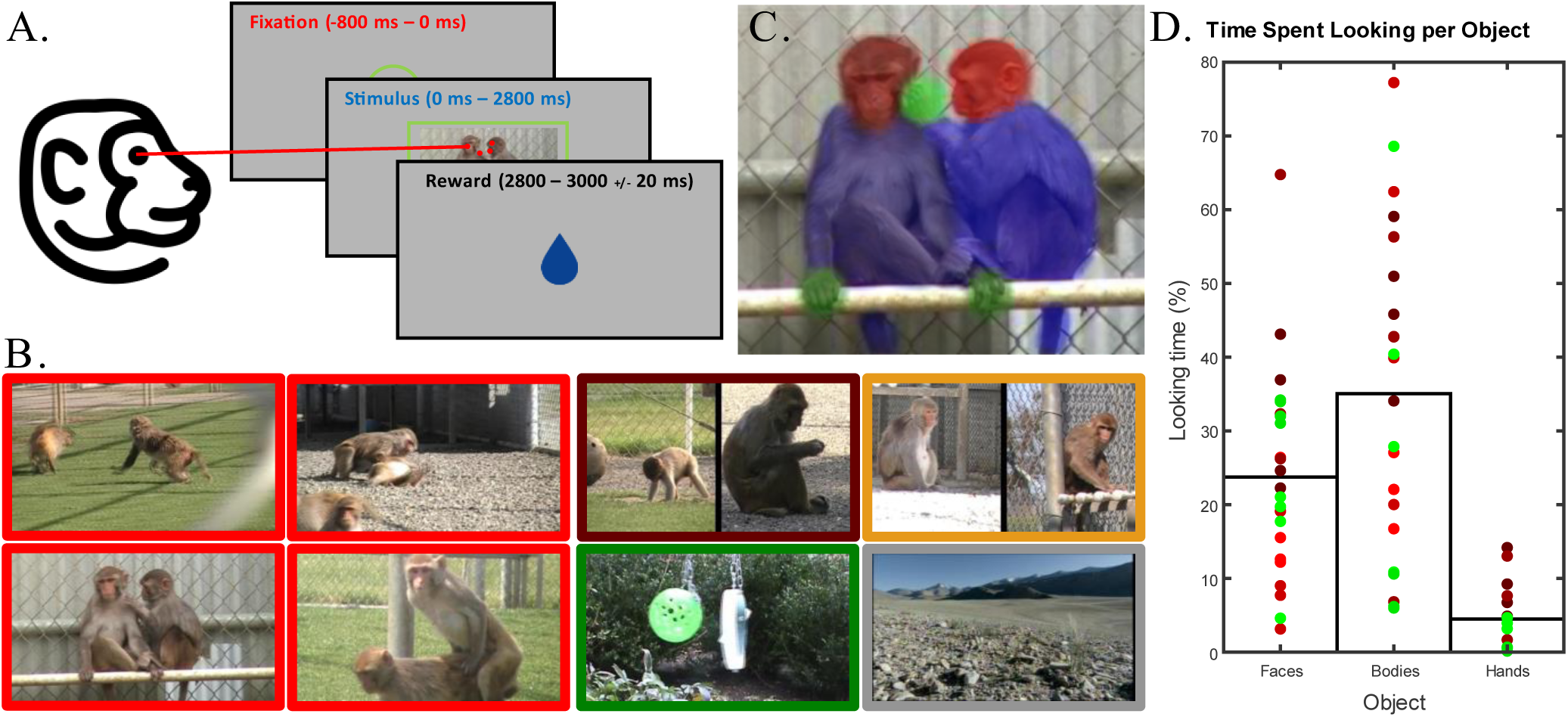
**A**: The Task Paradigm. 950 – 800 ms of Fixation is required to initiate trials, followed by 2800 ms of stimulus. Following successful free viewing of a box containing the stimulus and the surrounding 2 Dva, a 180 – 220 ms delay precedes a fluid reward. **B.** Trials consist of 2.8 second clips of videos containing either instances of chasing, fighting, grooming, or mounting conspecifics (social Interactions) or instances of two separate conspecifics engaging in goal directed behavior, Idling, as well as clips of two objects interacting or scenes without objects (non-social controls). **C.** Video contents were labeled to allow for tracking of subject’s gaze target – faces (red), bodies (blue), and hands (green). Hands were not labeled in chasing and fighting videos. **D**. Subjects actively scanned stimuli, spending the majority of viewing time focused on conspecifics, when present.

### Units responded to different phases of the task

We recorded from 1078 neurons in left-hemispheric medio-dorsal prefrontal area F7 in two macaque monkeys (*Macaca mulatta*). The target region was identified by structural MRI (see Methods). We found neurons responding during different periods of a trial. Some neurons responded transiently at fixation or video onset; in other neurons activity increased in a gradual ‘ramp’ during video presentation, and yet others during reward delivery (Fig. 2B). Out of a total of 1078 recorded neurons, 359 responded to the fixation phase of the task, with 171 significantly increasing and 188 significantly decreasing their activity (Fig. 2B, see Methods). 397 neurons responded during the early stimulus period (defined as 0 to 500 ms after stimulus onset): 250 up- and 147 downregulating activity. 445 cells responded during the subsequent “late” response period (500-2800 ms after stimulus onset), with 275 upregulating and 170 downregulating activity. During the reward period, a total of 366 neurons responded, with 242 upregulating and 124 downregulating activity. Thus, area F7 responds strongly to external events including the appearance of a fixation spot, video presentation, and the delivery of juice reward.

**Figure 2 –.**
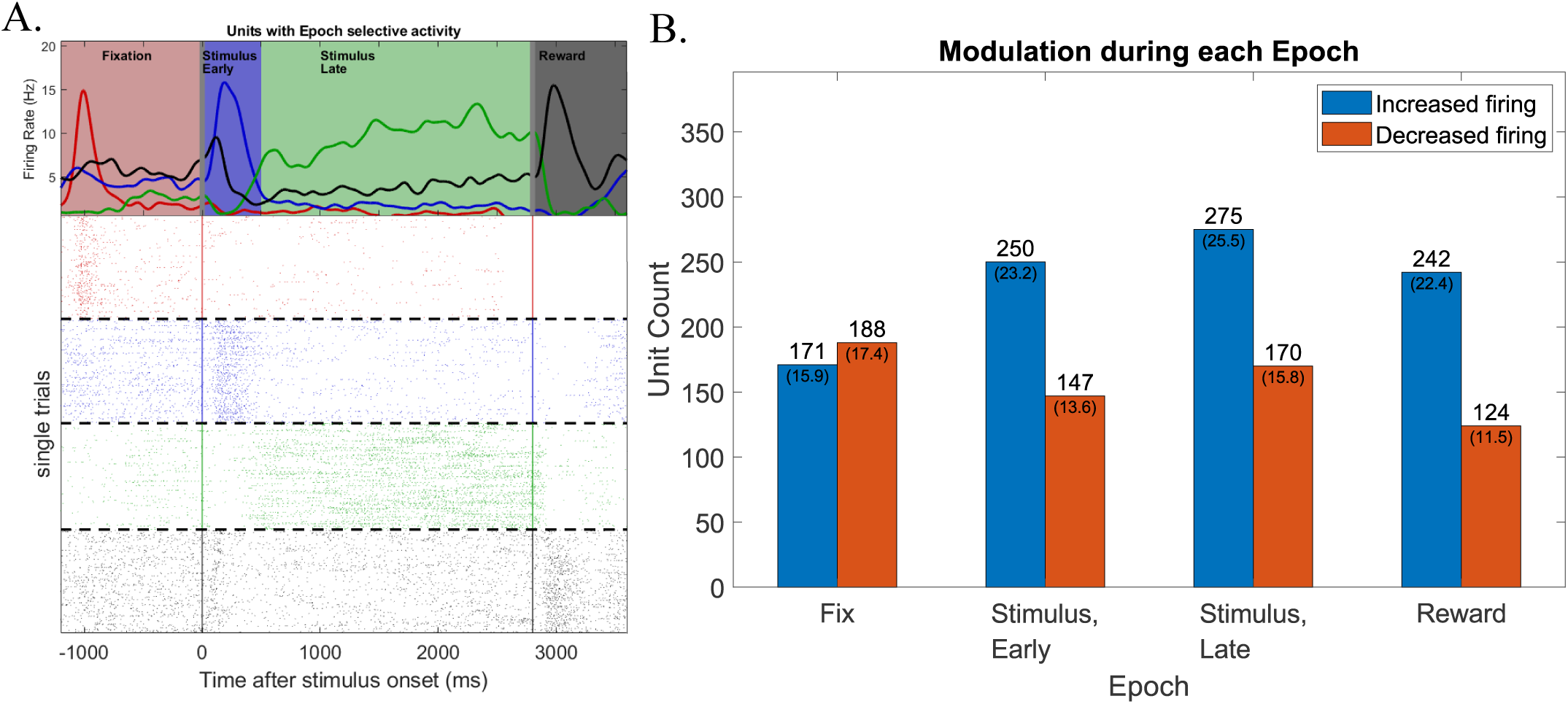
Activity is segregated across epoch lines. **A**: Example single unit rasters and Peristimulus time histograms of units with activity predominantly in a particular epoch of the task. **B**. Histograms illustrating total units modulated during each epoch of the task. 60.8% of units (652/1072) show some modulation from baseline during at least one epoch.

### Units with stimulus activity can differentiate stimulus content

A large fraction of the neurons in our sample (49.6%) responded to videos, and single unit responses varied across videos. Fig. 3 shows three example neurons that responded during video presentation. The cell in Fig 3A responded with an initial burst to most videos, then sustained its response most to a few videos containing social interactions. The cell in Fig 3B had a strong onset response to nearly all agent-containing stimuli and nearly no response to non-agent-containing stimuli and responded periodically to some agent-containing stimuli, while another example unit responded most strongly at onset to videos without agents (Fig 3C).

**Figure 3 –.**
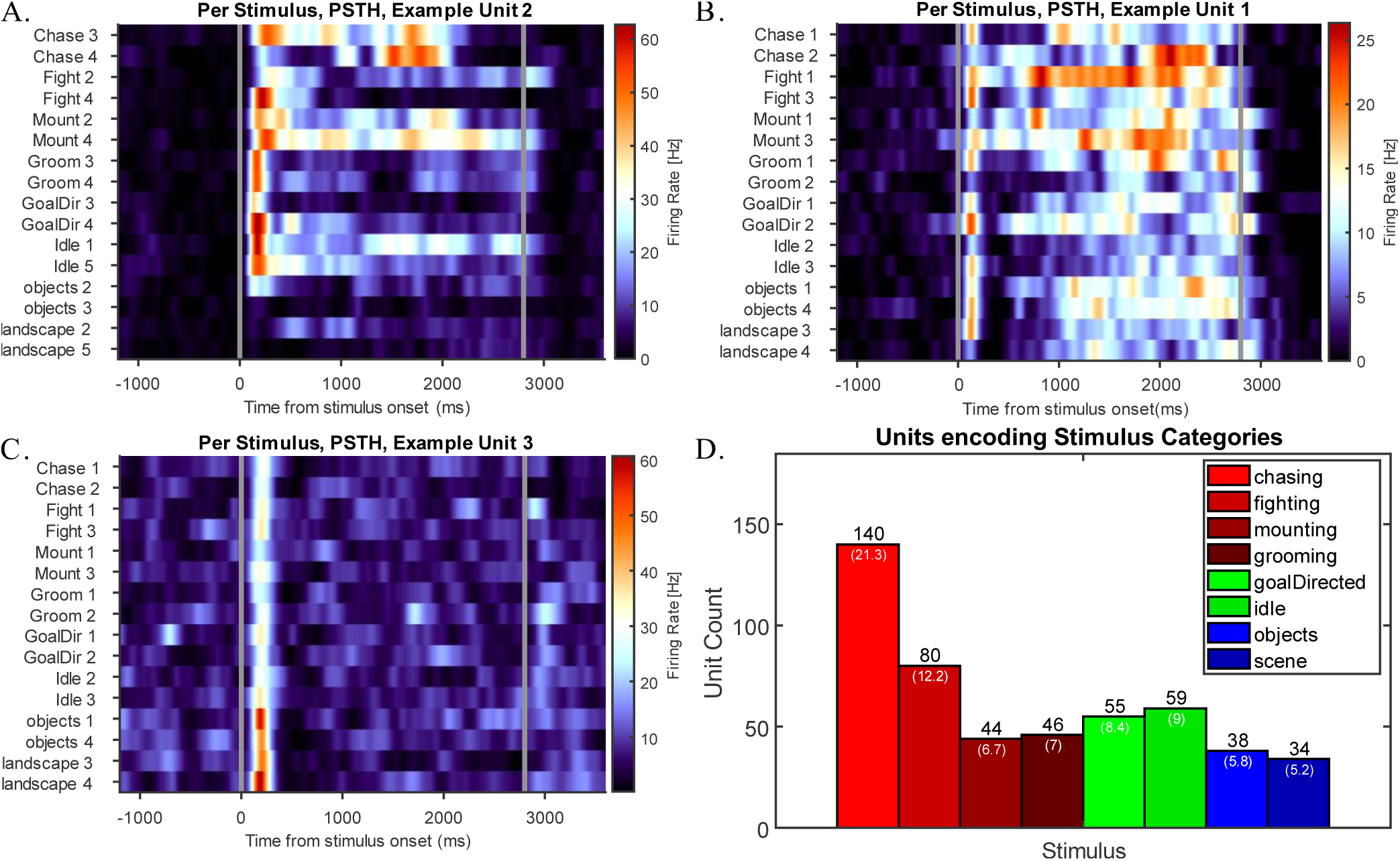
Units displaying different degrees of selectivity to categories. **A.** a unit displaying initial activity to most stimuli, but stronger activity to different instances of stimuli containing social conspecifics. **B.** a unit displaying initial activity to all agent containing stimuli, but more prolonged elevation activity to 2 instances of socializing conspecifics. **C**. a unit displaying strongest stimulus onset to all stimuli, with the strongest activity to stimuli lacking agents. **D.** a histogram displaying the number of units with significant stretches of activity for a particular category (see methods).

We first determined whether a neuron exhibited a significant response difference across stimulus conditions. Because such response differences could potentially occur at any time during a video, we measured this with a sliding window analysis (150ms bin size, 25ms step size). Units which showed a significant activity difference for a specific stimulus category for at least five consecutive bins, as assessed by an ANOVA, were classified as stimulus selective (see Methods). 432 neurons (40%) in our sample responded selectively by that definition (Fig 3D). Selective response enhancements were found for all types of video content: social interactions, non-social goal-directed behavior, idle behavior, objects, and landscapes. Thus, the population of area F7 cells exhibits diverse selectivities that may allow it to encode a wide variety of social and non-social scenes.

Given these selective response enhancements, we speculated that a population code may be present in area F7. We formally tested this possibility with a pseudo-population response decoding analysis (Meyers et al., 2015, see Methods). We found that the population response could indeed decode the different video categories (depicted in Fig 1B) on single trials with an accuracy of 67% across 8 categories (12.5% chance level). This decoding analysis used the time-average firing rate during stimulus presentation; high category separation was also visible in a time-resolved decoding analysis (300 ms bins with 100 ms steps) with on average 47.7% decoding performance across all categories (Fig. 4C). Decoding performance peaked at about 500 ms into the video presentation at ~70%, and then plateaued at ~45% for the remainder of video presentation. All stimulus categories were significantly decodable throughout presentation, but some much more so than others: chasing and the fighting categories were particularly decodable with average decoding of 90% and 73%, respectively. Thus the population of area F7 neurons encodes high level social and non-social variables across a wide variety of videos.

**Figure 4 –.**
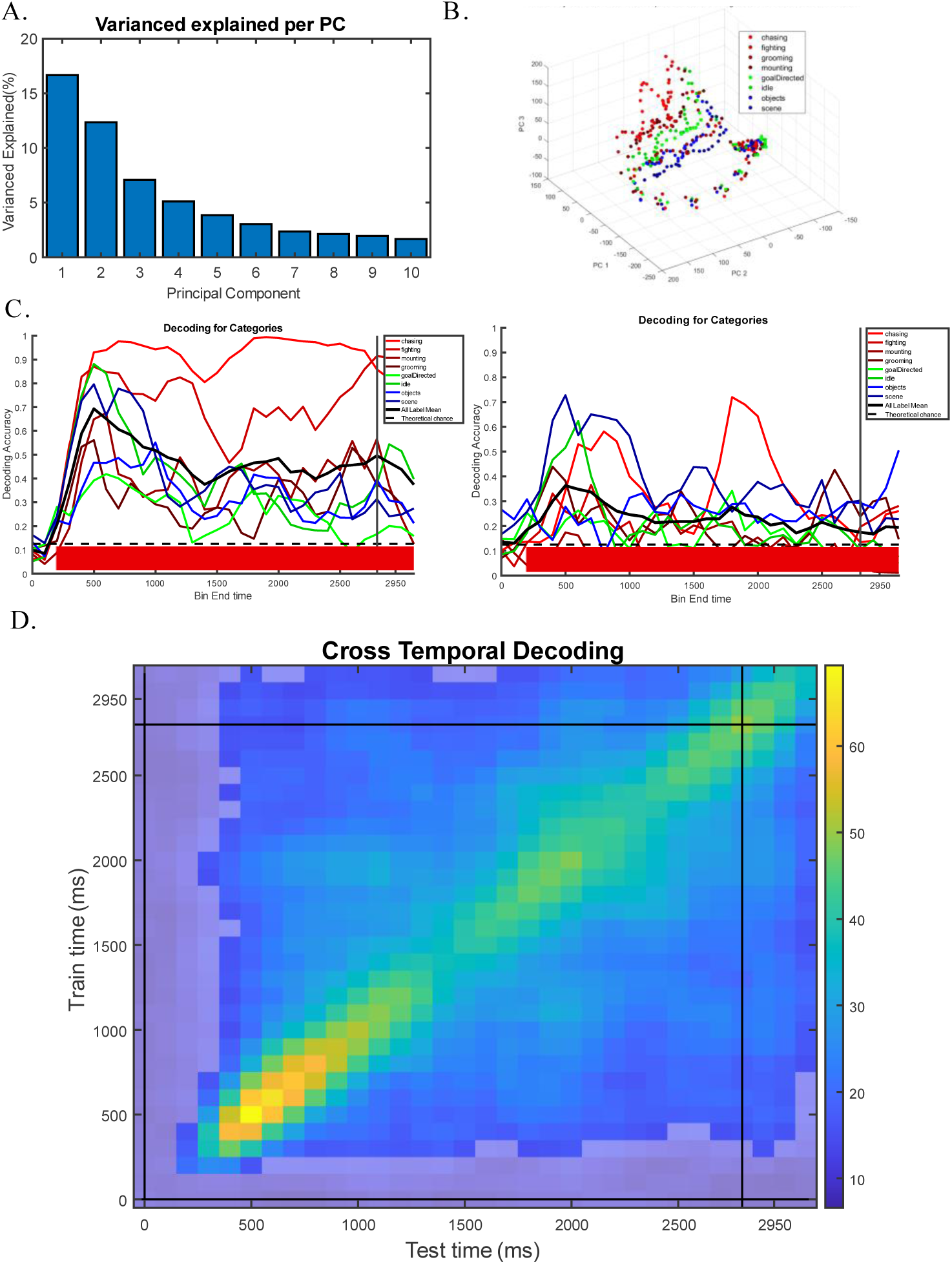
Units contained abstract category information. **A.** Principal components analysis performed on unit activity revealed a low dimensional space in which the top 10 principal components explain 51% of the observed variance in activity. **B.** The top 3 principal components explained 32% of the variance in the population. Activity occupies the same region of space for the duration of the fixation and reward periods, diverging during the stimulus presentation. **C. Left -** Maximum correlation coefficient decoding performed using a pseudo-population consisting of all recordings was able to successfully decode category with 40– 65% accuracy across stimulus presentation period, on average. **Right -** When trials for a specific stimulus are used only for testing or training of the model, decoding accuracy drops but remains significant for the duration of the trial. **D.** Cross temporal decoding, where B represent the diagonal. Dimed regions represent non-significant values. Though the strongest values reside on the diagonal, off-diagonal significant decoding suggest some stable coding.

To better understand the nature of the population code, we performed three further analyses. First, given that different neurons make different contributions to the population response throughout the duration of the video (e.g. Figs. 2A, 3A), we wondered how stable the code was over time. We performed a cross-temporal decoding analysis (Meyers et al., 2015) to determine how stable the population code was across stimulus presentation, i.e. while detailed content changed, but the overall meaning of the video remained the same (e.g. the details of a fight were changing, but the entire video showed one particular fight). To perform this analysis, we trained our classifier on data from one time point, and tested it using data across all time points (see methods). Fig. 4D shows decoding performance is lower off from the diagonal, indicating a strong dependence of the population code on moment-by-moment content. It also shows substantial consistency of the code after the early response period and throughout the late response period.

Results so far are based on individual videos and the populations ability to encode them. However, videos within each stimulus set were not randomly selected, but fell into different categories: two videos showed fighting, two chasing, etc. (see Methods). Thus, second, we wondered whether decoding might generalize across videos of the same category, suggesting that the population code carries abstract information about a chase going on or a fight. We performed a generalized decoding analysis in which we trained our classifier on one stimulus belonging to a category label tested it on the remaining stimulus. Decoding performance for this generalized categorical decoding (Fig. 4C, Bottom) dropped markedly relative to the stimulus agnostic one (Fig. 4C, top). Yet average categorical codes were significantly above chance (23%, Chance = 12.5%) throughout the duration of video presentation. Thus, similar to results of a cross-temporal decoding analysis, the population code of area F7 neurons depends substantially on detailed content, but also reflects categorical stimulus information.

Third, we sought to characterize the space of population responses. In particular we were interested to see if there was any response similarity structure reflecting high-level video content: decoding results would suggest that responses to within-category video should cluster, but there may be a larger overall structure of similarity, e.g. from social interaction to action to idle to non-agent containing video. To further characterize the population response, we performed a principal component analysis (PCA). We found that the population response was high-dimensional, with just over half (51%) of the variance carried by 10 dimensions (Fig. 4A). Some principal components differentiated between stimulus conditions (Fig. 4B). Within the sub-space of the top three principal components, which together captured ~31% of the variance, social interaction and non-social conditions were fairly separated (Fig. 4B). Thus the population response exhibits some systematicity separating high-level video conditions.

### Some units carry information about transient visual events

During the duration of video presentation, categorical content stayed constant, yet details changed. The aforementioned analyses point to area F7 neurons being sensitive not only to high-level video content, but also to more transient features. Yet, these analyses leave open the question as to which lower-level features might be driving neural responses. We performed an event-analysis of videos to isolate the occurrence of specific events like head turns, body turns, or ‘eye contacts’ (Fig. 5A, see Methods). Some of these events were only a few hundred milliseconds long, while others lasted much longer. Given that some cells responded only very briefly during some videos, we reasoned that it might be possible that they responded specifically to events and not only or not at all to the more abstract categorical content of the video.

**Figure 5 –.**
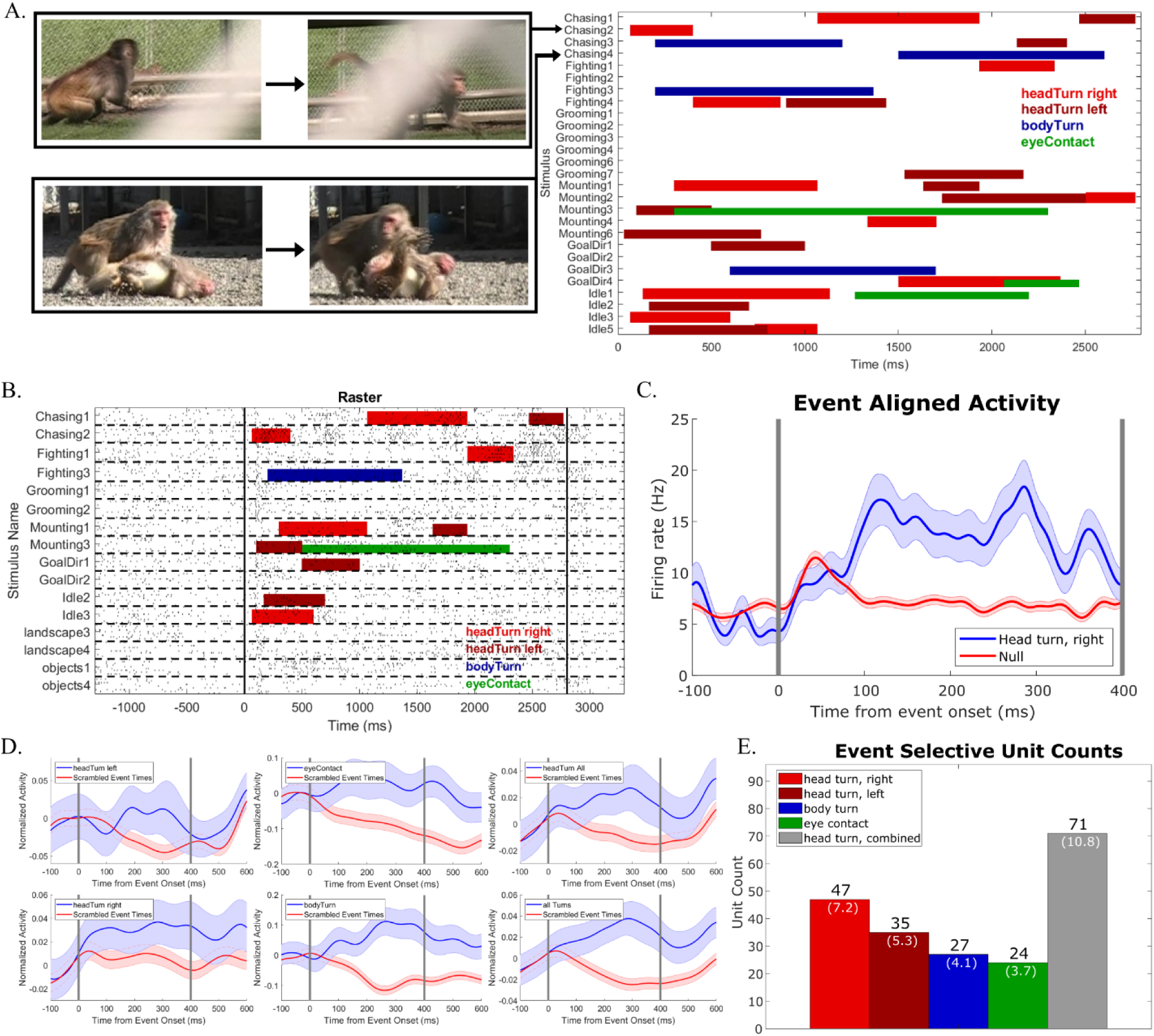
Head turning selectivity in the population of recorded units. **A.** Example frames from stimuli containing the instance before and after a head turn to the right. **B.** Top, Raster of unit with selectivity for head turning to the right. Bottom, Peri-stimulus time histogram of unit activity aligned to events. **C.** Event aligned smoothed PSTH of example unit shown in A and B. **D.** Normalized whole population mean responses to individual and merged labeled events. **E.** Population counts for selectivities across each labeled stimulus event. Percentages represent percentages of task modulated population.

Fig. 5B shows an example cell that responded selectively to right head turns (see Methods): activity was significantly higher following the onset of the rightwards head turning event than during the same period in stimuli during which the event did not occur (see Methods). The effects of these events were significant even in the average population response. Head turns, body turns, and eye contact all caused elevated activity at the population level (Fig. 5D). The effects were significant in 71 units (10.8%) for either left or right head turns, 27(4.1%) for body turns, and 24 (3.7%) for eye contact (Fig. 5E). Thus the F7 population was sensitive to relatively low-level, but socially conspicuous events.

### Reward-Related Activity

Activity during the reward period could be due to actual reward delivery or the expectation of it. To differentiate between the two possibilities, in our design we included 10% of trials during which no reward was delivered. Comparing activity during trials with and without reward delivery, we only found a response when the reward was actually delivered. Fig. 6A shows the average normalized response of the population of cells significantly modulated during this time period (n=267). The population response, furthermore, was following, not preceding reward delivery in these units, as illustrated by the example in Figure 6B. Responses to reward delivery were fast, with the population response peaking around 60 ms after reward onset. Thus area F7 responds quickly to reward delivery. During a portion of recordings, we observed units with ramping activity toward the end of the stimulus presentation window. An example is seen in Figure 6C. Though this activity may be in anticipation of a reward, our task was not designed to disentangle activity specific to the end of the stimulus rather than the expectation of a reward.

**Figure 6 -.**
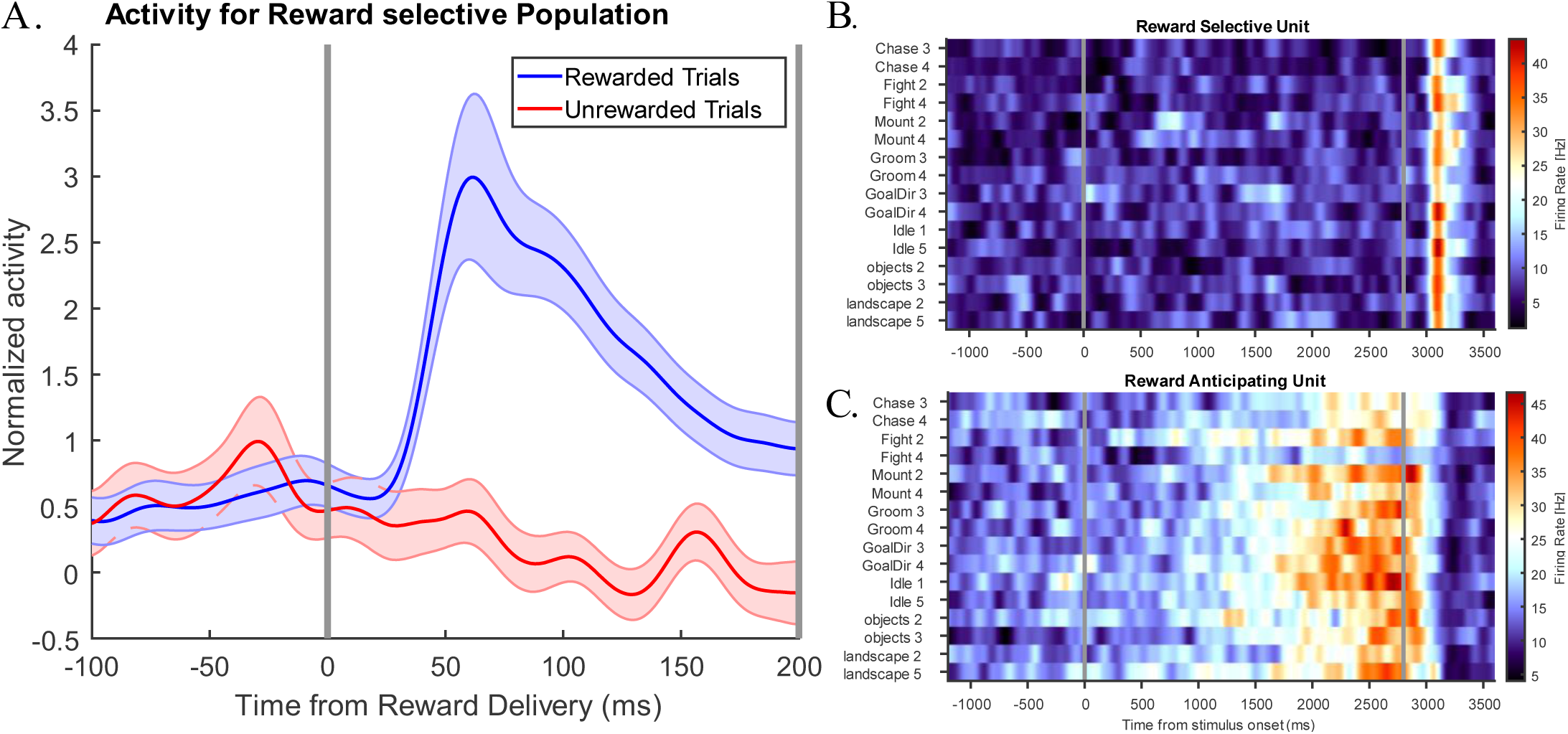
Reward related activity. **A.** Normalized activity across reward selective population of units, consisting of 267 units of 656 units. **B.** Example neuron showing reward selective activity. **C.** Example neuron showing ramping behavior at the end of the stimulus presentation period, terminated by the delivery of the reward.

### Sensitivity to the Subject’s Eye Movements

We analyzed the relationship of neural activity across many external stimuli including the fixation point, the videos, various specific events within them, and the reward stimulus. But during video presentation, subjects were free to explore the video through eye movements and thus to respond to the video. Fig. 7A shows a set of example movie frames and one subject’s eye movements during a set of trials. We wondered whether, in addition to all the aforementioned selectivity to external events, F7 neurons might carry information about the subject’s own actions. We focused on the subject’s saccades.

**Figure 7 –.**
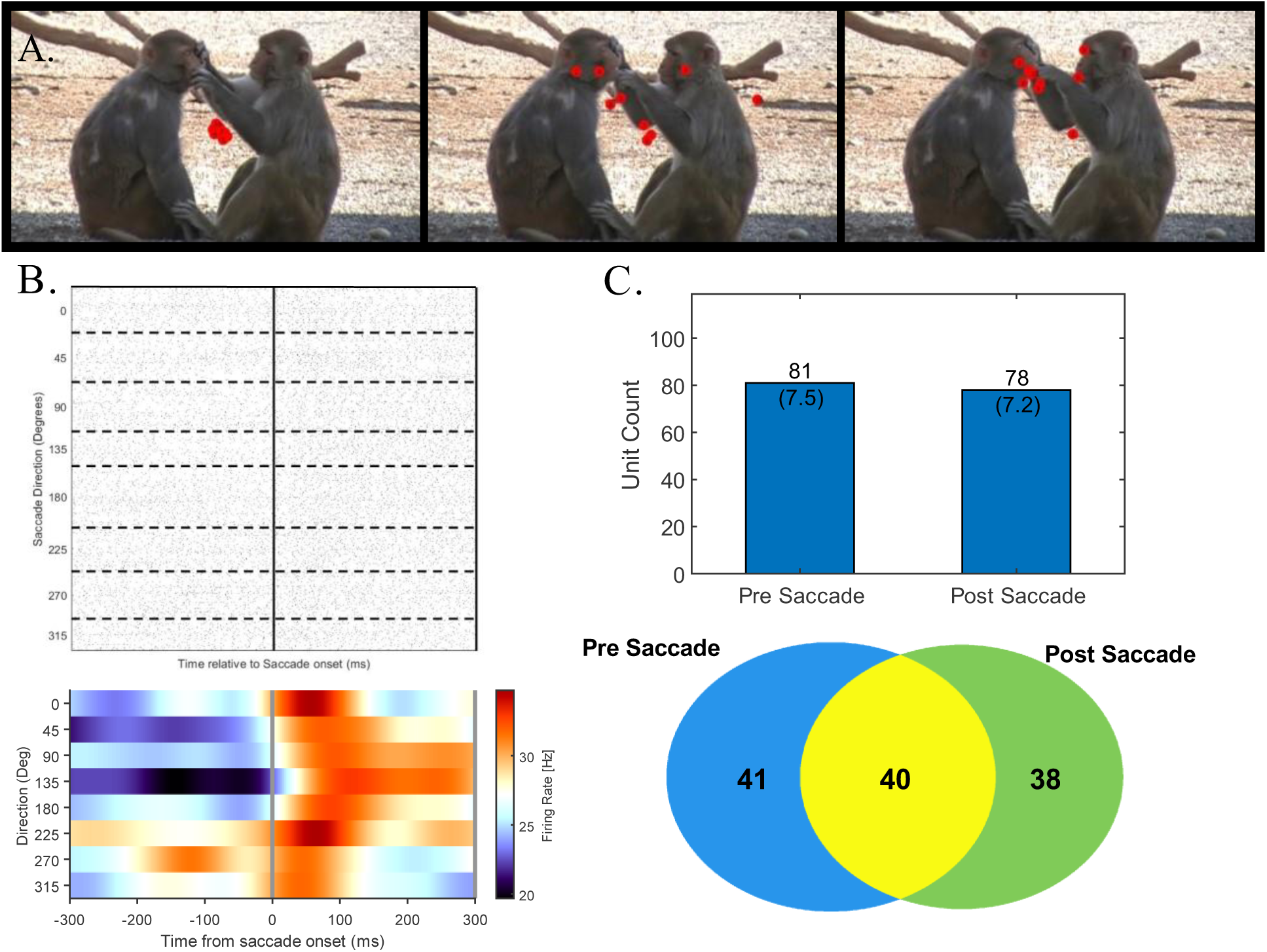
Saccade selectivity in the population of recorded units. **A.** Frames of stimuli with eye signal from different trials overlayed **B.** Top, Raster of unit with directional saccade selectivity, Bottom, Peri-stimulus time histogram of unit activity aligned to events. **C.** Bar plots representing number and percentage of units with pre and post saccade selectivity. **D.** Venn Diagram showing selectivity saccade activity across recorded units.

We detected saccades using the ClusterFix Toolbox (König & Buffalo, 2014), see Methods. We then binned activity in the pre- and post-saccade period and compared it to trials of the same stimulus where a saccade did not take place. We found many F7 neurons whose activity was modulated around the time of a saccade. Fig. 7B shows an example neuron whose firing rate was increased after a saccade in a direction-independent manner. We found a total of 81 neurons (18%) showing pre- and 78 neurons (17.3%) showing post-saccadic activity modulations (Fig. 6C), several times more than expected by chance (see Methods). Of these, 40 neurons showed both pre and post-saccadic activity (Fig. 6C, Bottom). Thus a substantial portion of neurons in 8B (26.9%) are modulated by the subject’s own actions in the form of saccades.

### Mixed selectivity across task and stimulus features

Recent work examining coding schema in the frontal lobes have posited neurons implemented a mixed coding scheme, to take advantage of the computational efficiency of high-dimensional representations (Fusi et al., 2016). Given the diversity of sensory and task variables represented by region F7, we wanted to determine whether neurons implemented a mixed coding scheme. Using previously obtained frequencies for each individual selectivity, we determined whether overlap across selectivities would be expected by chance.

Given how different the nature of the three external stimuli and of the associated responses (fixation, visual exploration, and awaiting reward), we wondered whether distinct or overlapping populations responded to the three different signals. We found that neural populations responding during fixation, videos, and the reward period largely overlap in a variety of combinations (Fig. 8A). Out of 1078 recorded neurons, the largest group sensitive to a period of the task, 172 neurons (16%), responded to all three periods of the task, 260 (24.1%) responded to two periods, and 224 (20.7%) responded to only one of the three event types. These values are higher than expected by chance (Fig 8A, lower left), many standard deviations above co-occurrence frequencies seen in scrambles (See Methods).

**Figure 8 –.**
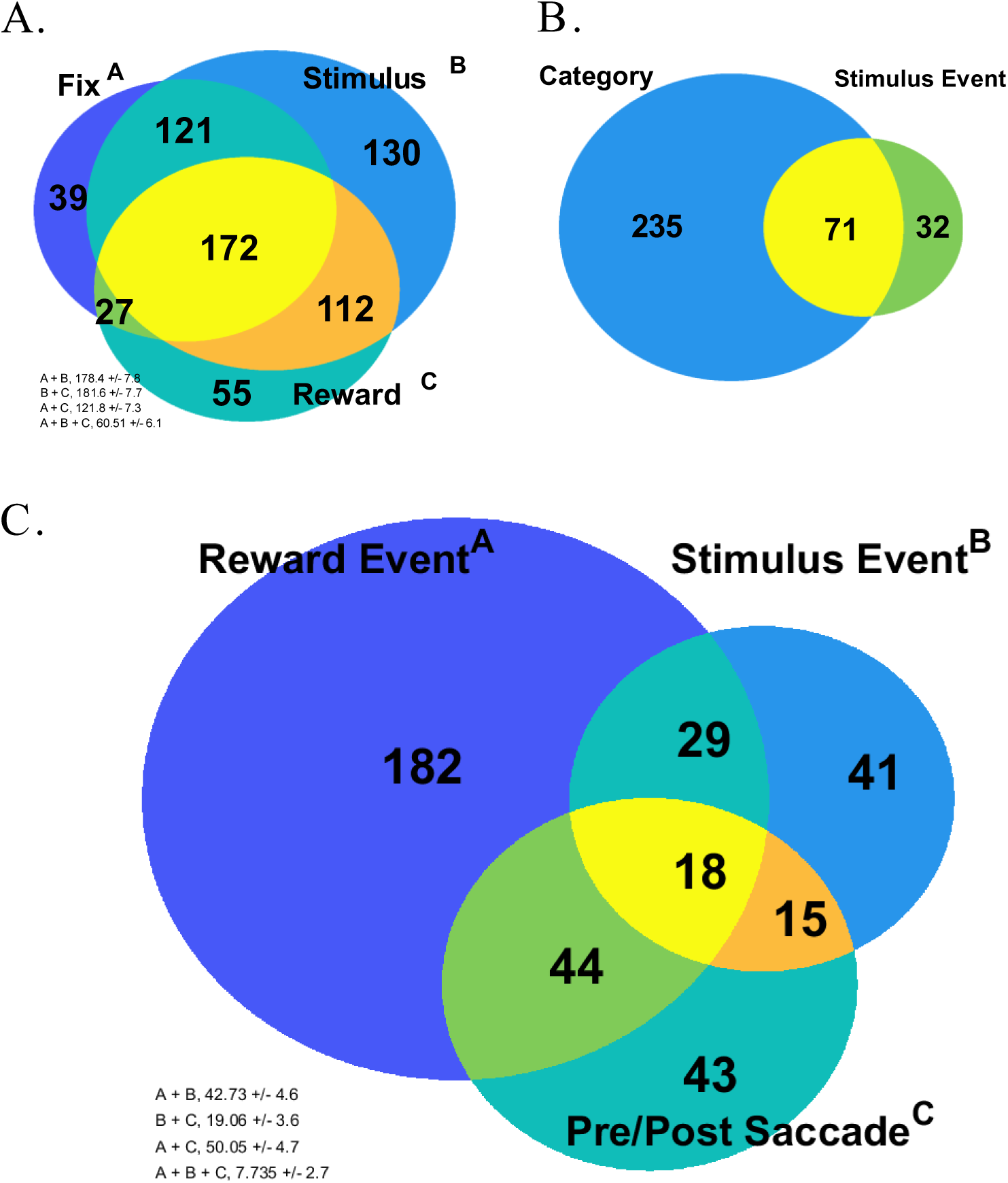
Mixed selectivity across recorded population. **A.** Overlap across populations of units modulated by different periods of the task. Expected overlaps calculated through simulation on entire recorded unit population (see Methods). **B.** Overlap across populations modulated by category, as determined by sliding window method, and units identified as event selective. **C.** Overlap across populations selective for the event of reward delivery, labeled stimulus events, and units modulated by the peri-saccade period. Expected overlaps calculated through simulation on entire recorded unit population (see Methods).

We next wondered whether different populations carried information for category than those that carried information for detail. We found though that these populations were overlapping (Fig. 8B). Of the units showing selectivity for an event of head turning, body turning, or eye contact, 69% (71/103) carried information about a category. Given the event selectivity may be sufficient to drive category selectivity in some instances, overlap is expected, though the fact some units which do differentiate across categories without representing labeled events suggests more abstract encoding may be taking place.

Given the multitude of mixed selectivity in area F7 neurons, we wondered whether these motor-related signals would be carried by neurons different from those responding to sensory events or would also be mixed. Of the units showing saccadic modulation, we found that the majority of units (64%, 77/120), this activity co-existed with selectivity for category, reward, or both. These values are close to those expected by random combination of feature selectivity across neurons (Fig. 8C).

## Discussion

In this study, we performed the first systematic exploration of dorsomedial frontal area F7 with social and non-social video. Combining this approach with multi-site recordings of a large population of neurons allowed us to explore how populations of area F7 neurons encode visual, social cognitive, reward, and motor behaviors – and thus whether a single function can be ascribed to this area. The results show that this one area encodes all of these functional levels, and a single functional role fails to capture this diversity.

Two main lines of evidence invoke area F7 as a core region of the primate social brain. First, area F7 was among a network of medial, dorsomedial, and ventrolateral prefrontal areas activated selectively while monkeys were watching video of the social interactions between other monkeys (Sliwa & Freiwald, 2017). It was therefore thought that these areas would support high-level social cognition, like the inference about the mental, emotional, and intentional states of other social agents or the elaboration, storage of socio-emotional scripts. Prior work supporting this role demonstrates strong functional connectivity between F7 and the temporal face patch system (Schwiedrzik et al., 2015). Thus area F7 is firmly established as part of the social brain and has been thought, prior to our study, to support high-level cognition. Yet what information it encoded and how was not directly known. Our electrophysiological recordings from large populations of F7 neurons sought to answer these questions. Results on both what variables the population encoded and how it accomplished this were highly surprising in light of current thinking about high-level social cognition. These findings have potentially far-reaching implications for the understanding and the study of frontal cortex and its relationship to sensory information processing at large.

The idea that area F7 extracts abstract information about social interactions and possibly infers the stable underlying, yet not directly observable mental states that generate them, predicts that when presented with video stimuli containing social interactions, the population of F7 neurons should carry information generalizing across the specifics of the interaction and across its evolution in time. We indeed found that the area F7 population did both (Fig. 4C). Furthermore, the F7 population response exhibited some systematicity in encoding high-level social content (Fig. 4D), which is also broadly compatible with the idea of abstract social content encoding.

Yet we also found evidence for encoding of concrete visual events within the video that was not necessarily social in nature and did not remain stable throughout: many area F7 neurons responded transiently to changes in visual stimuli, be they social or not, even just the onset of a fixation point (Fig. 2A) or video onset (Figs. 2A, 3A-C); and many F7 neurons responded to specific visual events like head turning, body turning, or eye contact, which are of potential social relevance (Fig. 5B-D). Given how many neurons responded to these events, it is possible that many area F7 neurons might encode yet other events. Furthermore, the population code was substantially more informative about specific video content than about social interaction categories, and showed as much evidence for changes over time as for stability (Figs. 4B-D). Maybe equally surprising as the richness of visual information encoded by area F7, was how fast responses could be triggered, in many cases in less than a hundred milliseconds. Thus in many ways area F7 neurons would be difficult to distinguish from neurons throughout the visual system. Thus area F7, beyond representing high-level social information, provides a rich representation of dynamic visual scenes.

This was possible for us to discover because we presented high-dimensional stimuli, natural video. This is not typical for the study of dorsomedial frontal cortex and, in fact, of frontal cortex at large. Past studies used low-dimensional paradigms in which subjects, typically after extensive training, were asked to interact with others and predict their actions (Chang et al., 2013; Haroush & Williams, 2015). Neural populations, similarly, were then analyzed with a question of whether these low-dimensional abstract concepts of others or rules were represented. Here we find that a region in dorsomedial frontal cortex contains a surprisingly high-dimensional representation of a real-world stimulus (Fig. 4A). Similar investigations in other cortical regions might similarly reveal that in addition to high-level and low-dimensional descriptions, rich and dynamic representations of stimuli might exist, possibly even, like here, of stimuli that are not even task-relevant.

## Methods

### Subjects and surgical procedures

Data were obtained from two male rhesus macaques (*Macaca multata*, 8 – 10 years old, 12 – 14 Kg). Standard anesthetic, aseptic, and postoperative treatment protocols were followed to implant MRI-compatible ceramic screws and acrylic cement, in which an Ultem head post was embedded for the purposes of head restraining. Subjects were trained to maintain fixation to initiate trials and to freely move their gaze around the stimuli using scrambles. Successful maintenance of fixation was rewarded with water. Following successful completion of 90% of trials, custom recording chambers were implanted, and craniotomies were performed to grant access to above anatomically localized area F7 (Saleem and Logothetis, 2012). All procedures conformed to federal and state regulations and followed the *NIH Guide for Care and Use of Laboratory Animals*. Surgeries and electrophysiological experiments were conducted at the Rockefeller University, and anatomical brain imaging was carried out at the Citigroup Biomedical Imaging Center (CBIC) at Weill Cornell Medicine. All experiments conducted were approved by the Institutional Animal Care and Use Committees (IACUC) of the Rockefeller University and Weill Cornell Medicine.

### Electrophysiology Recordings

Custom designed grids were printed in biocompatible Visijet M3 Crystal (*3D Systems, Rock Hill, South Carolina*). Grids were designed with 1 mm spacing between holes. Trajectories were targeted to the medial most region between the beginning of the principal sulcus and the end of the super limb of the arcuate sulcus. These trajectories were confirmed using recording grids, where tracks of interest were loaded with a polyimide tube containing a thin steel wire, visible on high resolution MRI scans of each subject. During recordings, 16 or 32 channel S-probes (*Plexon Inc, Dallas, TX)* were loaded into metal guide tubes, then positioned using 3D printed grids against the exposed and thinned dura. Electrodes were advanced using the Alpha EPS GII system (*Alpha Omega, Nazareth, Israel*). Each recording day, probes were advanced into the region until some channels showed distinct waveforms. Neural activity was amplified and sampled at 30 kHz by the Cerebus Neural Processing system, (*Blackrock Microsystems, Salt Lake City, Utah*). Extracellular waveforms were extracted and automatically clustered offline using the *waveClus 3* software package (Chaure et al., 2018). Units were manually curated, wherein results were checked and identified units unlikely to represent isolated units were unsorted.

### Behavioral Tasks and Visual Stimuli

Subjects were trained to complete trials which contained a fixation phase, a stimulus presentation phase, and a reward phase. Subjects had to maintain fixation on a centered spot (0.5 degrees of visual angle (dva)) within a circular window 4 dva in diameter for 800 – 950 ms (fixation phase), followed by free viewing of a rectangular space 14 * 22 dva for 2.8 seconds (stimulus phase), which contained the stimulus and an extra 2 dva on all sides. Upon successful maintenance of gaze within the defined regions for the duration of the fixation and stimulus phases, a juice reward was delivered after a 180 – 220 ms, where fixation was not required. During recordings, subjects were head-fixed inside a darkened Faraday cage, and visual stimuli were presented on a CRT monitor (NEC FE21111SB, eye-screen distance 62 cm, 35.65 * 26.73 dva) at a 60 Hz refresh rate. A photodiode placed at the lower right corner of the monitor was used to signal transitions in task phases. Eye position was recorded at 120 Hz using the ETL-200 Primate Eye Tracking System (*ISCAN, Woburn, Mass.*). The photodiode traces and eye data were collected, along with neural activity, by the Blackrock data acquisition system. Stimulus presentation and behavioral control were managed by the NIMH MonkeyLogic package, Version 2.2.2 (CITE)) run in MATLAB 2021a. This same version of MATLAB was used to perform all analyses.

### Stimulus Set

The entire stimulus library consisted of 32 color videos, 512 by 288 pixels in size, each shown for 83 frames at 30 frames per second. These stimuli were used in a prior study (Sliwa & Freiwald, 2017). These 32 color videos were divided into two sets (16 videos in each) balanced across categories. Each set contained the following:

- 8 videos of conspecifics participating in chasing, fighting, grooming, or mounting (two per category).
- 2 videos of conspecifics engaging in idling and 2 videos of conspecifics engaging in goal-directed behavior, such as manipulating a toy or eating a piece of food – These videos had been generated by concatenating 2 videos (512 * 140 pixels each) with a thin black bar down the middle.
- 2 videos of landscapes
- 2 videos of common objects – cage toys – suspended and colliding into each other.

#### Event Labeling

Stimuli contained brief movements or acts (‘events’, henceforth). Stimuli were watched and labeled when 4 distinct events took place – head turning to the left, head turning to the right, body turning in either direction, and eye contact. Eye contact specifies instances in the stimuli where a conspecific looks directly at the recording device, giving the impression of eye contact with the viewer.

### Data Analysis

#### Neuronal responses

Only trials during which subject maintained fixation successfully for the fixation period (the first 800 – 950 ms of the trial) and the stimulus presentation period (the subsequent 2800 ms) were used in the analysis. Stimulus-aligned spike trains were binned at 1 ms intervals and averaged across presentations of the same stimulus to create peri-stimulus time histograms (PSTHs). These were then smoothed with a gaussian kernel (σ = 40 ms) to estimate the spike density function (SDF). The following definitions were used to classify units:

##### Task Modulation

A unit which showed a significant increase or decrease in response to a particular task phase compared to baseline. Spike counts for each unit were taken per trial during the baseline (−1300 - −800), fixation (−800 – −1 ms), stimulus early (0 – 500 ms), stimulus late (501 ms – 2800 ms), and reward (2800 – 3200 ms) periods, and compared with a Mann Whitney U test, as implemented by the MATLAB function *ranksum*. Units which showed significant difference from baseline for at least one epoch were labeled task modulated.

##### Category Selectivity

A unit which showed a significant change in activity between categories during the stimulus presentation video. Activity was binned in 150 ms segments, incremented at 25 ms for each trial, and labeled according to the category of the stimulus. There were 8 possible categories – chasing, fighting, grooming, mounting, goal directed, idle, objects, and scene. Bins across these 8 labels were compared via an one-way ANOVA (MATLAB function *anovan*). Bins which varied significantly across labels (alpha = 0.05) were stored, along with the category label which differed. Units showing at least 5 consecutive bins of such selective activity for the same category were labeled category selective.

##### Event Selectivity

A unit which showed a significant change in activity after an event in the stimulus. Spike counts were taken in the 3200 ms following every instance of each event. Null periods were established by taking spike counts within the same 200 ms window of trials during which other stimuli were shown in which no event was taking place. Samples were compared using a Mann Whitney U test (MATLAB function *ranksum,* alpha = 0.05). Units in which spike counts significantly differed across stimuli with and without events in the aforementioned periods were labeled event selective.

##### Saccade Selectivity

A unit which showed a significant change in activity in the pre- (−200 ms to 0) or post (0 to 100 ms) saccade period. Saccade times were determined using the ClusterFix package (König & Buffalo, 2014). Resulting saccades shorter than 35 ms in duration or lower than 1 dva in distance were removed. Saccades times for saccades taking place outside of the stimulus presentation period were collected. Spike counts before (pre-saccade) or after (post saccade) were binned and compared to null bins. Null periods were established by taking spike counts within the same window during other trials where no saccade was taking place. Samples were compared using a Mann Whitney U test (MATLAB function *ranksum,* alpha = 0.05).

#### Population Overlap

Overlap between different labels was established through permutation testing. Each test involved generating a population matching the original in size, and randomly labeling units as selective for a particular label at the same frequency observed in the real population (i.e. 20% of the simulated units would be labeled as ‘event X selective’ at random in the simulation). Overlap between every combination of labels was determined for each of these simulated populations. This procedure was repeated 1000 times. The mean and standard deviation of the overlap seen across the 1000 repetitions between each label combination was reported.

#### Dimensionality Reduction

Principal components analysis was performed on population responses to gain an intuition for how population activity evolve overtime in response to different categories. Only units exceeding 1 Hz during the entire length of the average trial were included. Activity was then binned into non-overlapping 200 ms windows. Unit activity was averaged to all stimuli belonging to a particular category and concatenated into a bins * units matrix, prior to being processed by the MATLAB *pca* function.

#### Decoding Analyses

To assess population information content, unit rasters were binned in 300 ms windows in 100 ms steps. Pseudo-populations were generated by combining units across recording sessions, and the sum of trials for each label was randomly broken into a training set (3/4^th^ the trials) and a test set (remaining 1/4^th^ of trials) and trained with a maximum correlation coefficient decoder, implemented using the *Neural Decoding Toolbox* package (Meyers, 2013) through MATLAB. The same procedure was repeated after scrambling the labels of the training set to produce a null distribution of results, which were pooled across all bins. Decoding was determined to be significant if it exceeded the 95^th^ percentile of the null distribution.

